# Mutagenesis Mapping of RNA Structures within the Foot-and-Mouth Disease Virus Genome Reveals Functional Elements Localised in the Polymerase (3D^pol^) Encoding Region

**DOI:** 10.1101/2021.01.04.425359

**Authors:** Lidia Lasecka-Dykes, Fiona Tulloch, Peter Simmonds, Garry A. Luke, Paolo Ribeca, Sarah Gold, Nick J. Knowles, Caroline F. Wright, Jemma Wadsworth, Mehreen Azhar, Donald P. King, Tobias J. Tuthill, Terry Jackson, Martin D. Ryan

## Abstract

**Abstract:** RNA structure plays a crucial role in the replication of positive sense RNA viruses and can form functional elements within the untranslated regions (UTRs) and the protein coding sequences (or open reading frames (ORFs)). While RNA structures in the UTRs of several picornaviruses have been functionally characterised, the roles of putative RNA structures predicted for the ORF remain largely undefined. Here we have undertaken a bioinformatic analysis of the foot-and-mouth disease virus (FMDV) genome and predicted the existence of 53 evolutionarily conserved RNA structures within the ORF. Forty-five (45) of these structures were located in the regions encoding the non-structural proteins (nsps). To investigate if the structures in the regions encoding the nsps are required for FMDV replication we used a mutagenesis method, CDLR mapping, where sequential coding segments were shuffled to minimise RNA secondary structures while preserving protein coding, native dinucleotide frequencies and codon usage. To examine the impact of these changes on replicative fitness, mutated sequences were inserted into an FMDV sub-genomic replicon. We found that three of the RNA structures, all at the 3’ termini of the FMDV ORF, were critical for replicon replication. Contrastingly, disruption of the other 42 conserved RNA structures that lie within the regions encoding the nsps had no effect on replicon replication, suggesting that these structures are not required for initiating translation or replication of viral RNA. Conserved RNA structures that are not essential for virus replication could provide ideal targets for the rational attenuation of a wide range of FMDV strains.

**Importance:** Some RNA structures formed by the genomes of RNA viruses are critical for viral replication. Our study shows that of 45 conserved RNA structures located within the regions of the foot-and-mouth disease virus (FMDV) genome that encode the non-structural proteins, only three are essential for replication of an FMDV sub-genomic replicon. Replicons replication is dependent on RNA translation and synthesis; thus, our results suggest that the three RNA structures are critical for either initiation of viral RNA translation and/or viral RNA synthesis. Although further studies are required to identify if the remaining 42 RNA structures have other roles in virus replication, they may provide targets for the rational large-scale attenuation of a wide range of FMDV strains. FMDV causes a highly contagious disease posing a constant threat to global livestock industries. Such weakened FMDV strains could be investigated as live-attenuated vaccines or could enhance biosecurity of conventional inactivated vaccine production.

## INTRODUCTION

The genomes of RNA viruses not only encode proteins but also contain non-templated functional elements in both the coding and untranslated regions (UTRs). These can be secondary or higher order RNA structures such as simple stem-loops or more complex structures which include pseudoknots and so-called kissing-loops that mediate long-range RNA-RNA interactions (1–12) . The function, shape and number of such RNA functional elements is often characteristic for a particular group of viruses, where they play important roles in processes such as the initiation of viral RNA translation and replication, subgenomic mRNA transcription, frame shift events, viral RNA encapsidation and modulation of host’s antiviral responses (reviewed in (13)). Since many RNA viruses are of medical and veterinary importance, characterisation of these RNA structures brings us closer to understanding viral pathogenicity and provides opportunities for disease control.

Foot-and-mouth disease virus (FMDV) is the causative agent of foot-and-mouth disease (FMD), a highly contagious disease of cloven-hoofed animals (including livestock) (reviewed in (14)). FMD is endemic in Africa and Asia, where it impacts upon productivity and trade as well as posing a constant threat of causing costly incursions into disease-free countries (15–21). Control of FMD by vaccination in endemic settings is complicated by the high antigenic variability of the seven serotypes of FMDV: A, Asia 1, C (not reported since 2004), O, Southern African Territories (SAT) 1, SAT 2 and SAT 3 (19, 20, 22–24).

FMDV is a small non-enveloped positive-sense single-stranded RNA virus classified in the species *Foot-and-mouth disease virus*, genus *Aphthovirus* in the family *Picornaviridae*. The FMDV genome is ~ 8.5 Kb in size and composes of a single, long open reading frame (ORF) which is flanked by 5’ and 3’ UTRs (reviewed in (25)). The encoded polyprotein is co- and post-translationally cleaved by viral proteases (L^pro^ and 3C^pro^) and by a ribosomal skipping event mediated by the 2A peptide into a number of functional precursors and the mature proteins (26–34). The coding sequence for the FMDV ORF is arbitrarily divided into four regions (5’-L^pro^, P1, P2 and P3-3’). The P1 region encodes the capsid proteins (1A, 1B, 1C and 1D, also called VP4, VP2, VP3 and VP1, respectively), while the P2 and P3 regions encode the non-structural proteins (nsps) (reviewed in (25)).

There are a number of RNA structures within picornavirus genomes that have been accurately predicted and characterised biochemically (12, 35–39). These structures are predominantly located in the UTRs and have been shown to be important for replication and translation of picornavirus genomes (reviewed in (40)). Within the 5’ UTR of the FMDV genome, the S-fragment forms a single, long hairpin structure (293-381 nucleotides (nts) in length) and has been reported to play a role in viral replication and innate immune modulation (41–45). Elsewhere in the 5’ UTR, the presence of multiple (2 - 4) pseudoknots downstream of the poly(C) tract has been shown to determine virus tropism (41, 46). Other key and well-characterised RNA structural elements include a type II internal ribosome entry site (IRES), which initiates cap-independent translation of the viral genome (41, 47–50); while the *cis*-acting replication element (*cre*) acts as a template for uridylylation of the VPg (3B) protein, which then acts as a primer for synthesis of viral RNA (51, 52). The 3’ UTR of the FMDV genome is located upstream of the poly(A) tract and contains two RNA stem-loop structures called SL1 and SL2. These stem-loops interact non-simultaneously with the S-fragment and IRES forming long-range interactions that have been shown to be necessary for viral RNA replication (43, 53, 54).

A number of other secondary RNA structures have been predicted computationally to be present within the FMDV ORF (12). However, with the exception of packaging signals (55), the role(s) of these structures in the FMDV replication cycle have not been determined. In this study we have identified 45 evolutionarily conserved RNA structures within the regions of the FMDV ORF that encode for the nsps. Mutagenesis of these structures identified three novel RNA stem-loops in the coding region of the RNA-dependent RNA polymerase (3D^pol^) that are essential for replication of an FMDV sub-genomic replicon, suggesting that these structures are required for either initiation of viral RNA translation and/or viral RNA synthesis. In contrast, mutagenesis of the remaining 42 structures had no effect on replicon replication. This approach can aid in the identification of critical viral RNA structures required for viral genome replication, and also help identify conserved RNA structures that are not essential for virus replication that could provide ideal targets for the rational attenuation of a wide range of FMDV strains.

## RESULTS

### Prediction of conserved RNA structures within the FMDV genome

While previous studies have provided evidence that the FMDV genome is highly structured with conserved RNA base pairing throughout the coding part of the genome (12, 56), these studies were conducted on a relatively small dataset. Since the number of full genome sequences available on public databases has greatly increased in recent years, before conducting functional studies, we revisited these analyses to predict conserved RNA stem-loops that were common in 118 representative genomic sequences covering all FMDV serotypes (see materials and methods section for isolates information).

Firstly, average mean folding energy differences (MFED) across whole FMDV genomes were determined for all viral isolates used in this study. In this method, conserved minimum free energy (MFE) values were normalised to MFE values of native sequences that had been scrambled using an NDR algorithm, which preserves the dinucleotide frequencies of native sequences. This ensures that reported values are not purely due to G+C or other composition biases (see material and methods for detail) (57–59). In order to show distribution of the MFED values along the genome, this analysis employs an incremental sliding window computation with user-defined window size and increment (60) (in our case 400 and 20 nts, respectively, where each 400 nts segment overlapped its neighbours by 380 nts). A 400 nts window allowed for detection of the S-fragment structure, while ignoring potential long-distance RNA-RNA interactions for which biological significance is hard to verify. Despite the high genomic sequence diversity across all seven serotypes (20% mean nucleotide pairwise distance (± 9% standard deviation (StDev)), with 31% (± 5% StDev) and 14% (± 7% StDev) average pairwise distance in the regions encoding the capsid proteins and the nsps, respectively), all the FMDV genomes analysed showed high folding energies across most of their sequence compared to the permuted controls (Fig. 1). This indicates that all FMDV sequences possess a similar extent of sequence order-dependent RNA secondary structure. To confirm this, full genome sequences were grouped into those of Eurasian (A, Asia 1, C and O serotypes) and SAT (SAT 1 - 3 serotypes) origin and average MFED values were determined along the genome for each group. Although we recognize that the grouping may not completely accommodate the inter-serotypic history of these viruses (see (45) for details why grouping viruses into SAT and non-SAT clusters is not always correct), the MFED plots showed similar patterns of high and low MFED values across the genome. MFED values were better correlated between FMDV groups in the UTRs and the regions encoding the nsps identifying a potentially greater degree of RNA structure conservation compared to the more genetically divergent region encoding the capsid proteins (Fig. 1).

**Figure 1:**
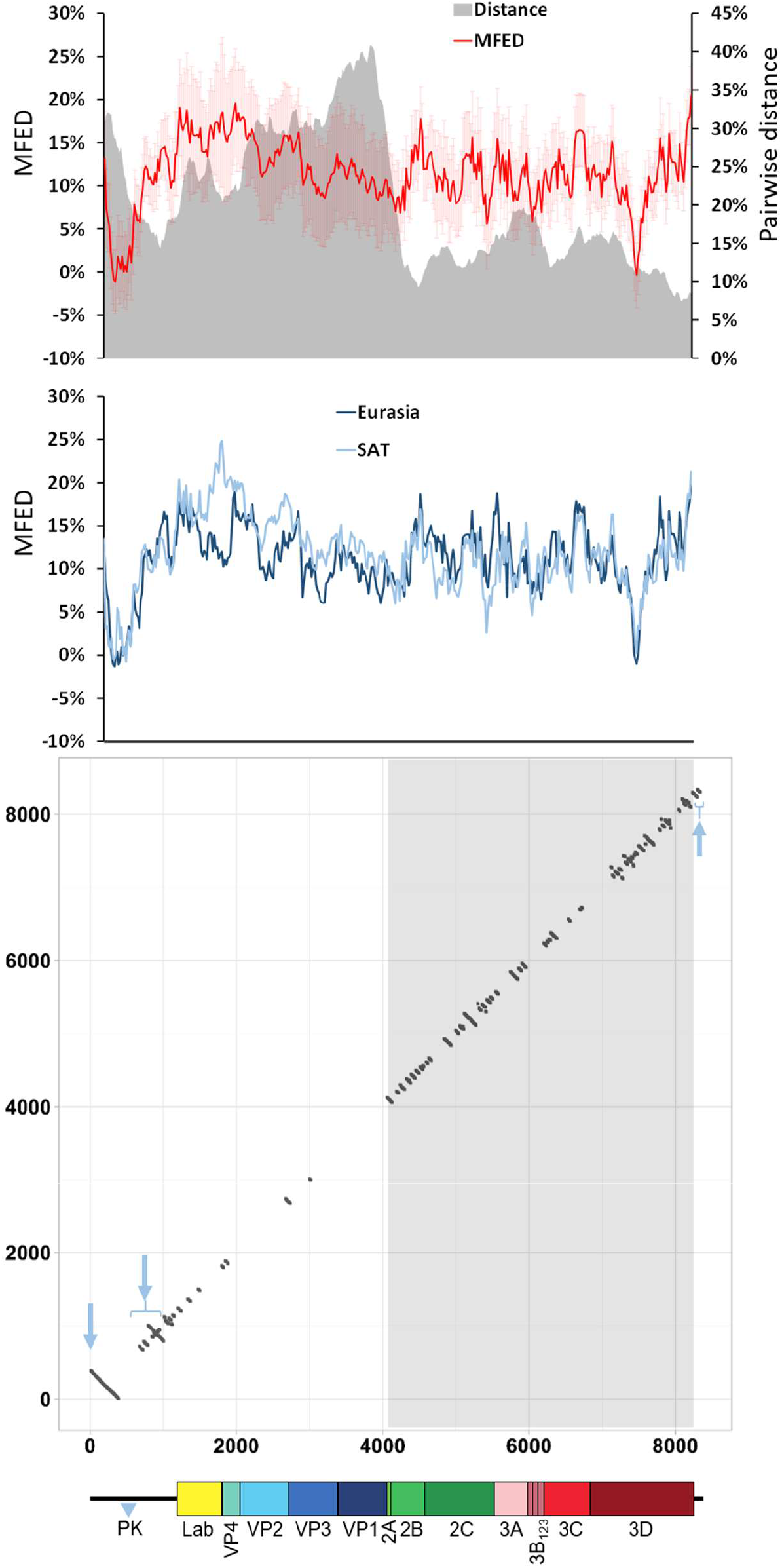
Extent of the conserved RNA secondary structures within FMDV genome. Upper panel shows a scan of pairwise distance and mean folding energies difference (MFED) prepared using SSE v1.4 software and 118 FMDV sequences representing all seven serotypes. The mean values for successive 400 nt fragments across the genome are plotted (where each 400 nts segments overlapped its neighbours by 380 nts). The light red shading represents error bars showing standard deviation from the mean for each datapoint. The middle panel shows MFED values for the same FMDV genomic sequences but grouped into Eurasian (A, Asia 1, C and O serotypes) or SAT (SAT 1 - 3 serotypes) clusters. The lower panel shows a dot plot graphical representation of RNA structures that were conserved across all seven FMDV serotypes. The x-axis and y-axis represent FMDV genome positions, with each dot representing a single pairing between two nucleotides, one with its position marked on the x-axis and the other one with its position marked on the y-axis. The three pale blue arrows indicate location of the S-fragment, *cre*+IRES and SL1+SL2 structures on the dot plot graph, respectively (for a detailed visualisation of these structures see Fig. S1). The blue tringle marked PK indicates the genomic region containing pseudoknot structures which was excluded from these analyses. The area corresponding to the regions encoding the non-structural proteins (i.e., P2 and P3) is highlighted in grey and for clarity, a schematic representation of the FMDV genome is drawn to scale.

The window size used for MFED scanning does not identify individual RNA structures and only highlights regions with high folding energies (which may contain dissimilar structures and/or structures located at different positions). Therefore, RNAalifold program, implemented in The ViennaRNA Package (61), was used to identify individual conserved RNA stem-loops for the 118 whole genomic sequences and for individual FMDV serotypes. Stem-loops that were conserved in all seven serotypes were visualised as a dot plot graph, plotting each nucleotide pairing (represented by individual dot) against positions of involved nucleotides on the x and y axes (Fig. 1). Any pairing interactions distanced by more than 400 nts were removed post analysis. By excluding long-distance interactions post whole genome RNA structure prediction, we did not ignore the effect they may have on formation of local pairings. RNAalifold cannot predict pseudoknots, and therefore the region directly downstream of the poly(C) tract was excluded from our analyses (Fig. 1).

These analyses correctly predicted the presence of well-characterised RNA secondary structures in the FMDV genome: the S-fragment, IRES and *cre*, all located in the 5’ UTR, and SL1 and SL2 located in the 3’ UTR (Fig. 1 and Fig. S1). It additionally identified several serotype-specific conserved stem-loops in the region encoding the capsid proteins, but only four of these were conserved in all seven serotypes. In contrast, 45 stem-loops (when counting each RNA hairpin individually, even within a single branched structure) were universally present within the regions encoding the nsps (Fig. 1, Table 1). Overall, there were 53 highly conserved stem-loops in the ORF of the FMDV genome that were conserved across all serotypes (Table 1).

**Table 1.** Number of conserved stem-loops within each FMDV genomic region

| Genomic region | Number of predicted stem-loops<sup>a</sup> |
| --- | --- |
| S-fragment | 1 |
| The rest of 5' UTR<sup>b</sup> | 11<sup>*</sup> |
| L<sup>pro</sup> | 4 |
| 1A (VP4) | 0 |
| 1B (VP2) | 1 |
| 1C (VP3) | 1 |
| 1 (VP1) | 2 |
| 2A | 1<sup>**</sup> |
| 2B | 7 |
| 2C | 10 |
| 3A | 3 |
| 3B<sub>1</sub> | 1 |
| 3B<sub>2</sub> | 2 |
| 3B<sub>3</sub> | 1<sup>***</sup> |
| 3C | 3 |
| 3D | 17 |
| 3' UTR | 2 |

<sup>a</sup>Each hairpin loop was counted individually;
<sup>b</sup>Excludes poly(C) tract and pseudoknot regions;
<sup>\*</sup>*cre* (a single hairpin loop), IRES domain 2 (a single hairpin loop), IRES domain 3 (five
hairpin loops), IRES domain 4 (two hairpin loops), IRES domain 5 (a single hairpin
loop), plus a single hairpin loop downstream of IRES;
<sup>\*\*</sup>Four nucleotides of the 5' end of the stem belong to the 1D encoding region;
<sup>\*\*\*</sup>17 nucleotides of the 5' end of the stem belong to the 3B<sub>3</sub> encoding region.

### Use of CDLR mutagenesis for functional mapping of predicted RNA structures

Next, we undertook mutagenesis studies to investigate whether any of the conserved RNA structures identified in the FMDV genome play a functional role in viral replication. FMDV replicons lack the region encoding the capsid proteins but are replication competent, demonstrating that there are no RNA elements essential for translation or replication of viral RNA within the capsid encoding region. Therefore, our investigation focused on structures located within the regions encoding the nsps of the replicon. Additionally, the effect on replication of changes incorporated into the replicon can be analysed in real-time through monitoring of fluorescence from an integrated green fluorescent protein (GFP) reporter gene that replaced the region encoding the capsid proteins (62).

In order to mutate the conserved RNA structures predicted within the regions encoding the nsps while maintaining codon composition, codon order and dinucleotide frequencies of the native WT replicon sequence we applied CDLR scrambling method (11, 56). To monitor its effectiveness in altering or otherwise disrupting RNA pairing within the native sequence, sequence of the regions encoding the nsps of WT replicon was randomly permutated 50 times using the CDLR algorithm. Then, MFED values for these mutants were calculated as described above and these were compared to MFED values of the native WT replicon sequence and the corresponding sequences of the 118 FMDV isolates used in this study. Sequences generated by CDLR showed evidence of severely disrupted RNA secondary structures, with a mean MFED value of 2.2% (StDev ±1.4%), compared to a mean value of 10.9% (StDev ±1.2%) for the corresponding regions of the native FMDV sequences and that of the WT FMDV replicon (Fig. 2).

**Figure 2.**
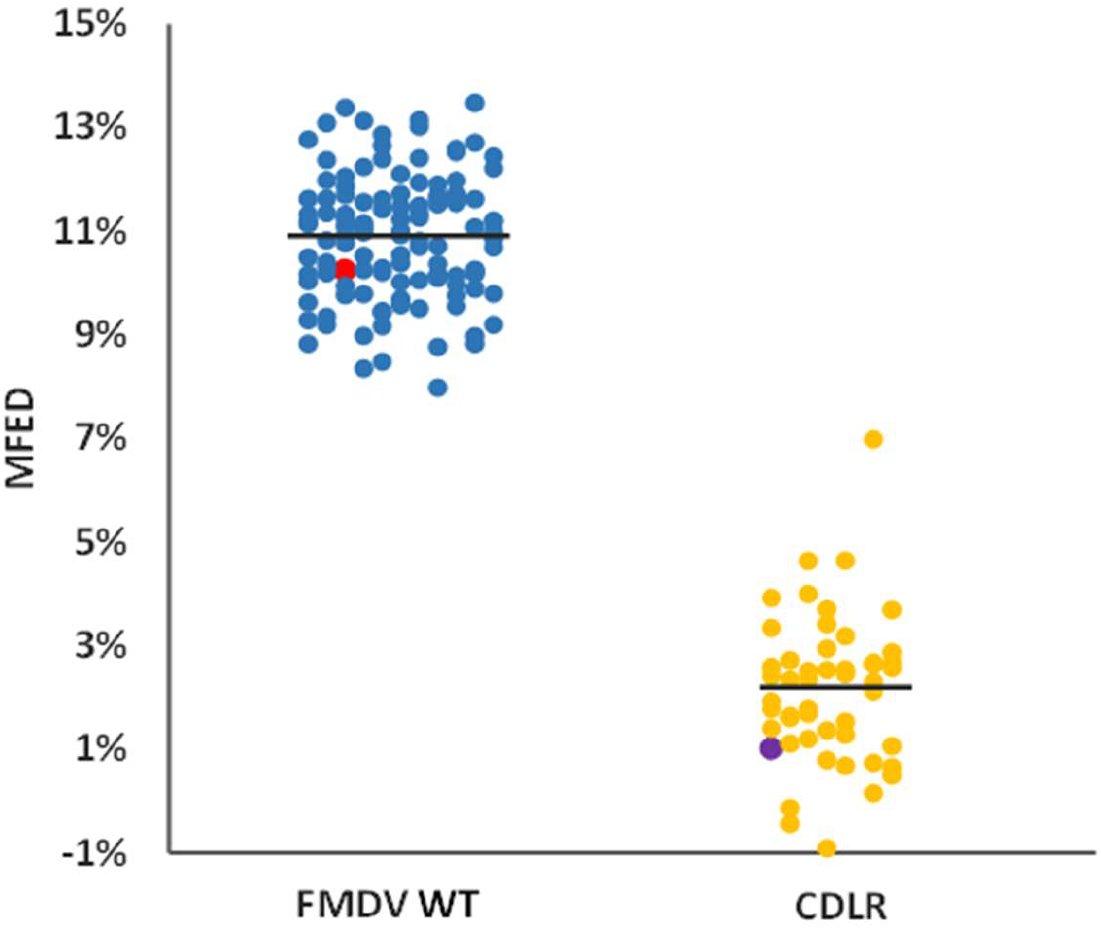
Comparison of average MFED values for wild type (WT) and CDLR-scrambled sequences. Mean folding energy difference (MFED) for the regions encoding non-structural proteins (nsps) of 118 FMDV field isolates representing all seven serotypes (blue dots), WT ptGFP-replicon used in this study (red dot) and CDLR scrambled sequences (yellow dots). Among the latter is the CDLR scrambled sequence used in this study to generate replicon mutants (purple dot). To obtain CDLR-scrabbled sequences the sequence of the regions encoding the nsps of the WT replicon was permuted 50 times by codon-shuffling to minimise RNA secondary structure, while preserving protein coding, native dinucleotide frequencies, and codon usage.

To identify functional RNA structures, we divided the regions encoding the nsps of the WT replicon (ptGFP-replicon) into nine consecutive fragments defined by unique restriction sites, and individually permutated each fragment using the CDLR algorithm (Fig. 3A-B). To further verify the extent of changes to the RNA structure introduced by the CDLR algorithm, we used the RNAforester program implemented in The ViennaRNA Package (61, 63, 64). This compared the putative structures adopted by the CDLR-permuted regions (shown in Fig. 3A-B) to the structures located within the corresponding regions of the WT replicon sequence. RNAforester calculates RNA secondary structure alignments based on the tree alignment model and quantifies similarity of structures in question, where the relative similarity score values equal to one represent two identical structures (61, 63, 64). With the exception of the 2C encoding region, which exhibits some structure similarity between CDLR and WT replicon (Fig. S2), there was low structural similarity between equivalent WT and CDLR genomic fragments (Table 2). RNA structures located in the 5’ and 3’ UTRs were generally unaffected by any CDLR permutation of the adjacent or more distal regions encoding the nsps, with the exception of the SL1 stem-loop in the 3’ UTR that was shorter by 11 pairings (Fig. S3).

**Table 2.**
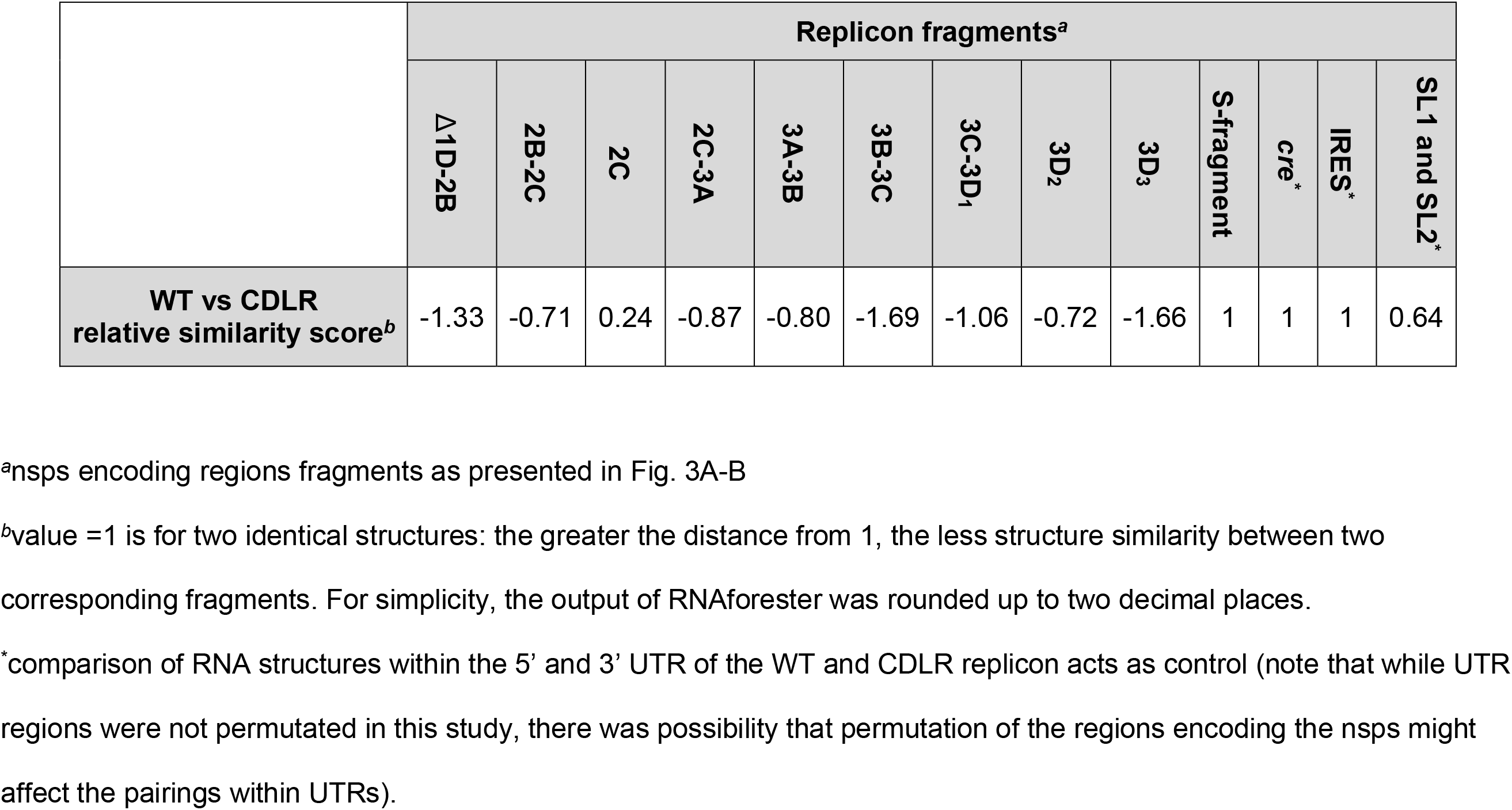
Similarity comparison of RNA structures within corresponding WT and CDLR replicon genomic fragments, calculated using RNAforester program

**Figure 3.**
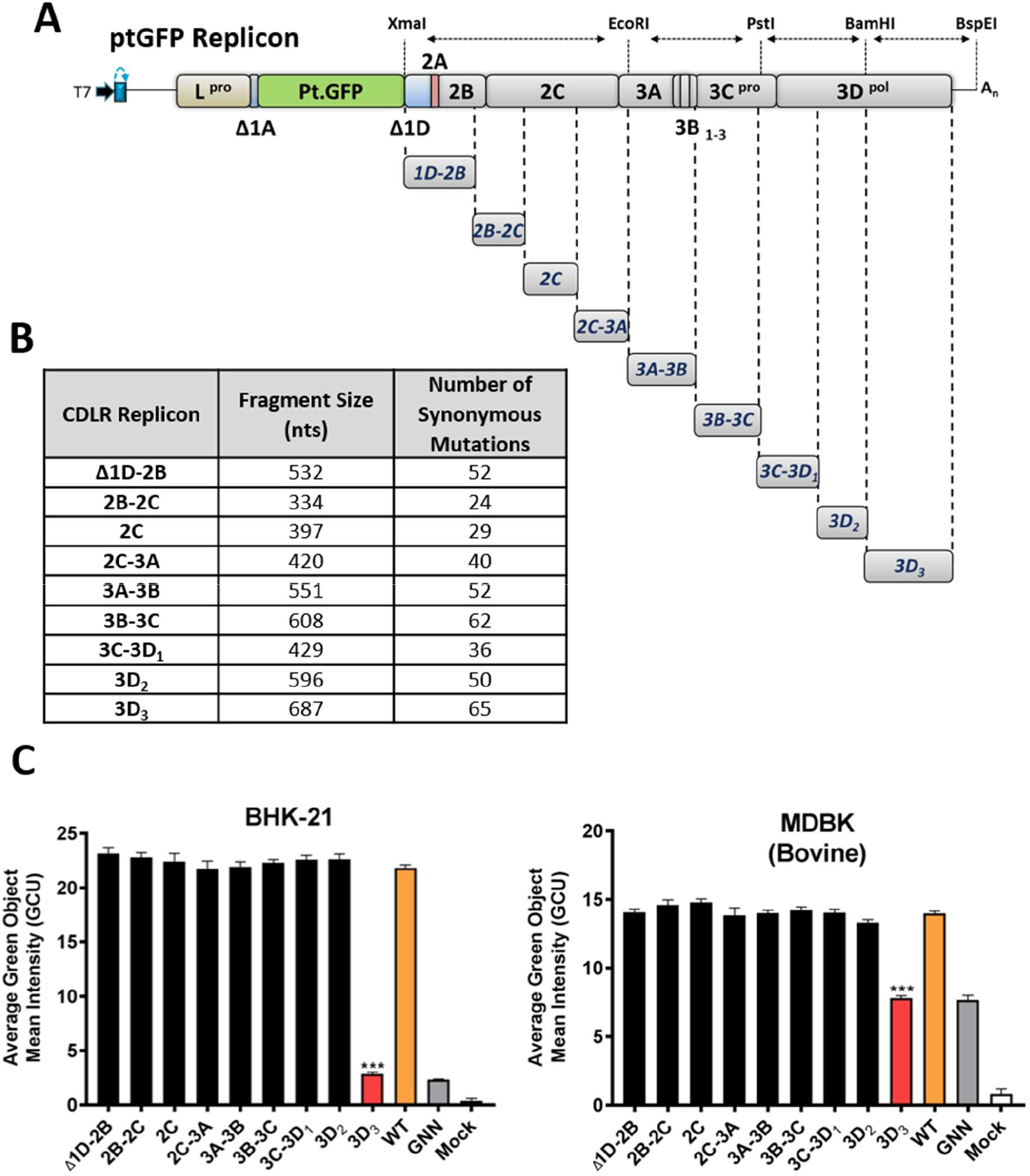
Replication of CDLR replicons within BHK-21 and MDBK cells. **(A)** Schematic representation of CDLR replicons. Mutated regions were firstly inserted into a sub-clone encoding the non-structural proteins (nsps) of the genome (Δ1D-polyA) before cloning into the WT ptGFP-replicon using the unique restriction enzymes shown. **(B)** CDLR replicon insert sizes and number of mutations within each region. Regions were chosen based on restriction site usage within the regions encoding nsps. Mutations were introduced as described within the materials and methods section. **(C)** IncuCyte data represent the average cell (green object) GFP intensity per well at 8 h post-transfection. Results are the mean of three independent experiments ± standard error. Significant differences between WT ptGFP and CDLR replicons were determined (***, P < 0.001). The replication-incompetent 3D^pol^ active site mutant (GDD → GNN) ptGFP-3D^pol^GNN was used as a negative control.

### CDLR replicon mutants reveal regions of secondary structure required for replication of an FMDV replicon

Next, we examined the effect of RNA structure disruption on replication of the FMDV replicon using mutant replicons containing CDLR-permuted sequences over different parts of the regions encoding the nsps. For this we used two different continuous cell lines known to support FMDV replication (Fig. 3A-B). The replication kinetics of the mutant replicons was compared to the WT ptGFP-replicon and a replicon with an inactive polymerase (ptGFP-3D^pol^GNN, previously described in (65)). Since replication levels at 8 hours post-transfection (hpt) were representative of the entire experiment (Fig. S4), for simplicity, data for this time point are shown. In both cell lines (BHK-21 and MDBK, of hamster and bovine origin, respectively), all of the CDLR mutant replicons tested displayed replication kinetics comparable to the WT ptGFP-replicon except for the replicon which carried a mutated sequence within the 3’ terminal part of the 3D^pol^ encoding region (called 3D_3,_ Fig. 3C). The replicon with 3D_3_ mutated encoding region was replication defective in both cell lines, with replication levels equivalent to the negative control replicon (ptGFP-3D^pol^GNN; Fig. 3C). These results strongly suggest that this part of the 3D^pol^ encoding region contains RNA structures crucial for replication of the FMDV replicon. Consistent with their inferred location in 3D_3_, CDLR permutation of the entire Δ1D-3A and 3A-3D_2_ encoding region showed little effect on the replication kinetics (Fig. S5).

### Modification of individual stem-loops within the 3D_3_ region impairs replication of an FMDV replicon

Our results indicate that the region of the FMDV genome encoding for the 3’ terminal end of 3D^pol^ (called here 3D_3_) contains conserved secondary RNA structures that may be necessary for replication of the FMDV replicon. Therefore, the RNA structures present in this region were investigated in more detail by visualising each individual structure and comparing it to the corresponding scrambled region within the CDLR mutant. Analysis of corresponding sequences of FMDV field isolates (over the 3D_3_ region) revealed five stem-loops (SL7 – SL11) with strong nucleotide pairing conservation, with SL10 being the most conserved structure (Fig. 4A). Variability within all structures was accommodated though the occurrence of covariant changes that preserved nucleotide pairings (Fig. 4A). Additionally, there was substantial nucleotide sequence conservation in the sequence forming the unpaired loop at the top of the stem-loop structures (i.e., in the hairpin loops) of SL7, SL8 and SL9 (Fig. 5) implying some functional constraints on these sequences. Each of the predicted structures in the WT sequence were substantially disrupted in the CDLR scrambled mutant (Fig. 4B).

**Figure 4.**
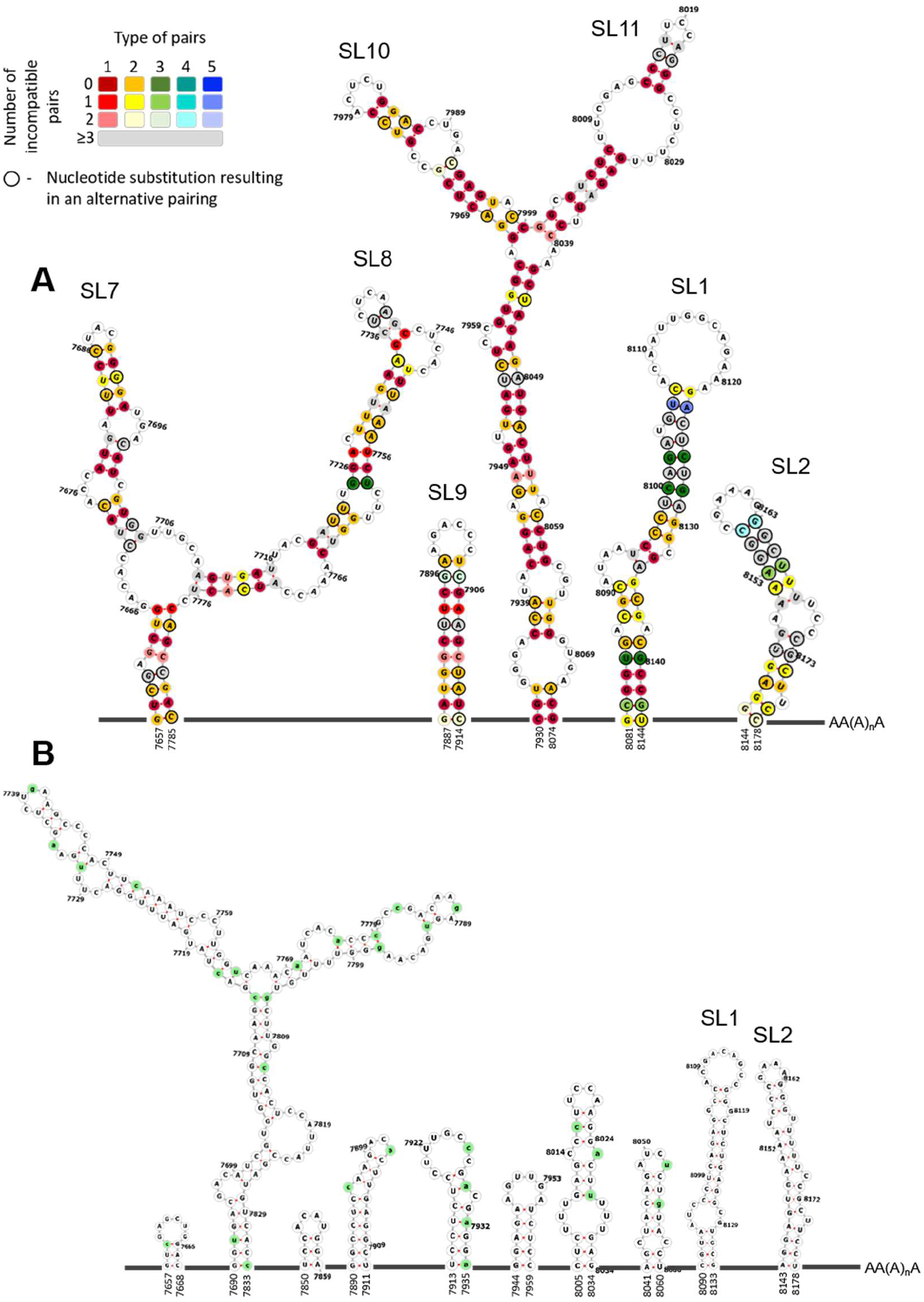
Schematic representation of predicted conserved RNA structures located at the 3’ terminal end of the region encoding 3D^pol^. **(A)** Schematic representation of conserved (in all FMDV serotypes) RNA secondary structures located at the 3’ terminal end of the region encoding 3D^pol^ (i.e., the 3D_3_ region described in Fig. 3). Conserved putative stem-loops (SL7 – SL11) are shown, where two stem-loops located in the 3’ UTR described before (SL1 and SL2) act as a control of the computational prediction. Nucleotide positions which form conserved pairing were colour-coded according to number of pairing types (‘red = 1’ to ‘blue = 5’) and conservation of a pairing (‘dark shades = nucleotide pairing occurred in all FMDV isolates’ to ‘light shades = lack of nucleotide pairing in two FMDV isolates’). Positions coloured in light grey show lack of pairing for three or more FMDV isolates. Black circular outline indicates nucleotide position where a substitution resulted in an alternative pairing (see included legend for detail). Unstructured regions are represented as dark grey lines and are not drawn to scale. Numbers represent nucleotide positions corresponding to the sequence of A/Brazil/1979 isolate (GenBank accession number AY593788). Supplementary Table S1 specifies details represented graphically in the figure legend. **(B)** Schematic representation of RNA secondary structures located in the 3D_3_ region after scrambling using the CDLR algorithm, demonstrating how RNA secondary structure in this region was changed. Mutated nucleotide positions are highlighted in green. Unstructured regions are represented as dark grey lines and are not drawn to scale. Numbers represent nucleotide positions corresponding to the sequence of the A/Brazil/1979 isolate.

**Figure 5.**
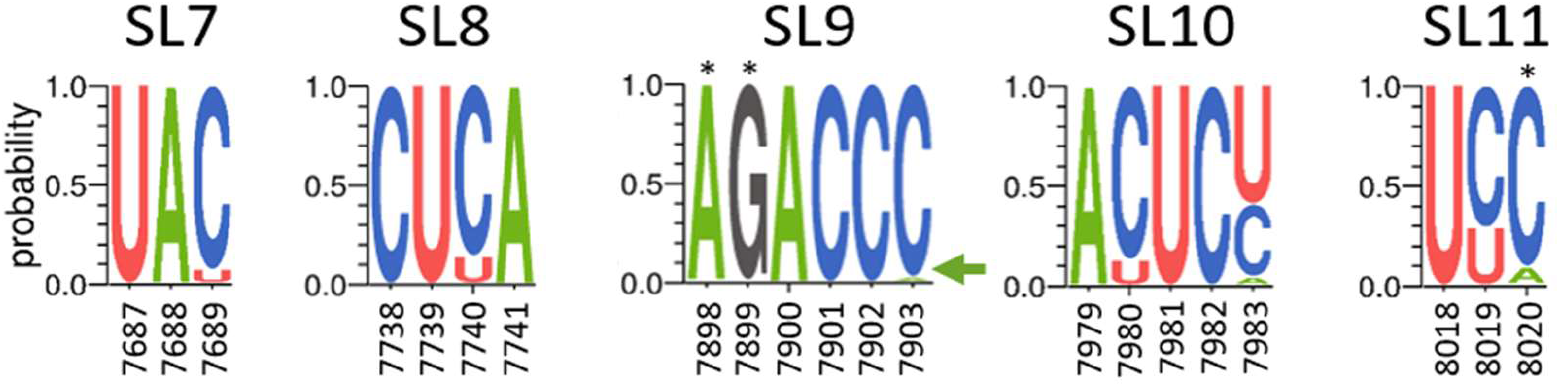
Extent of nucleotide conservation within hairpin loops of SL7 - SL11 RNA structures. Sequence logos were generated using WebLogo 3.7.4 web server based on sequences of 118 FMDV isolates. Probability shows the extent of nucleotide occurrence at a given position. Numbers represent nucleotide positions corresponding to the sequence of A/Brazil/1979 isolate (GenBank accession number AY593788). Asterix (*) marks positions where substitution occurs in 1 out of 118 FMDV isolates but due to a limited resolution of the y axis it does not appear in the sequence logos (these are: A7898G, G7899A and C8020G). The green arrow points to C7903A substitution which due to height of the A symbol could go unnoticed.

Further studies were therefore undertaken to dissect the importance of the individual stem-loops within the 3D_3_ fragment for replication of the FMDV replicon. Each of the five putative RNA structures in the 3D_3_ region of the WT replicon were permuted individually *in silico* introducing the maximum number of nucleotide changes possible to disrupt the RNA structure whilst maintaining amino acid coding, dinucleotide frequencies and the integrity of the neighbouring RNA structures (Fig. 6 and 7A). Additionally, a replicon where all five putative RNA stem-loops were altered (SL7-11^mut^, using the same mutation strategy as for each individual loop, Fig. 7A) acted as a negative control (in addition to the replicon with CDLR-scrambled 3D_3_ region) to confirm that mutation of these particular stem-loops, and not of other elements present in the CDLR replicon with the permutated 3D_3_ region, impaired RNA replication. Replication of ptGFP-replicons carrying individual mutated stem-loops was tested in the same two cell lines as described above (Fig. 7). As previously observed, replication levels at 8 hpt were representative of the replication kinetics (Fig. S6). Replication of replicons with disrupted SL7 and SL8 was not affected in either cell line (Fig. 7B). In contrast, replication of replicons with disrupted SL9 or SL10 were significantly reduced, although the effect on replication varied between the cell lines. Disruption of SL9 led to only a marginal, but statistically significant, reduction of replication in BHK-21 cells (GFP intensity equal 94% of the GFP signal of the WT replicon, p-value=0.02), whereas the negative effect on replication in MDBK cells was greater (GFP intensity equal 49% of the GFP signal of the WT replicon, p-value < 0.001). In both cell lines, disruption of SL10 reduced replication to a greater extent than disruption of SL9 (GFP intensity, 52% (p-value < 0.001) of the GFP signal of the WT replicon in BHK-21 cells, and 24% (p-value < 0.001) of the GFP signal of the WT replicon in MDBK cells), with the replication profile in bovine cells being close to the replicon with an inactive polymerase (ptGFP-3D^pol^GNN) and the replicon with the 3D_3_ region mutated by the CDLR algorithm (Fig. 7B). Replication of the replicon with disrupted SL11 was reduced only in MDBK cells (GFP intensity equal 85% of the GFP signal of the WT replicon, p-value < 0.001), but not BHK-21 cells. Finally, the replicon with all five stem-loops altered (SL7-11^mut^) demonstrated replication comparable to the ptGFP-3D^pol^GNN replication-deficient control (which give a GFP signal due to translation of the input RNA) in both cell lines tested (~20% of WT GFP signal, p-value < 0.001, Fig. 7B).

**Figure 6.**
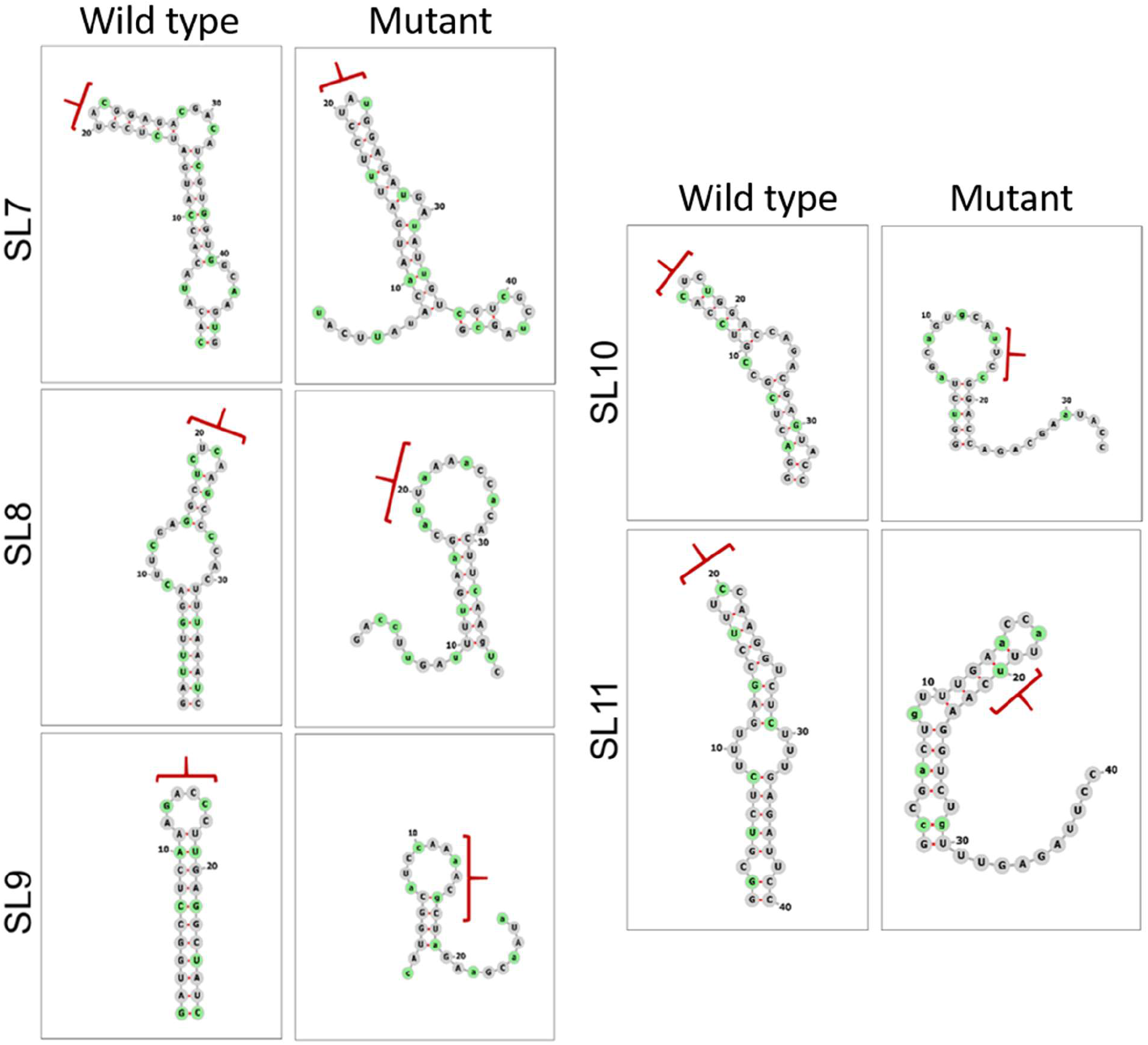
Disruption of the predicted RNA secondary structures by silent mutagenesis. The conserved stem-loops identified in the 3’ terminal end of the region encoding 3D^pol^ (i.e., 3D_3_) of FMDV were predicted individually by Mfold for the WT ptGFP-replicon. Predicted WT stem-loops were mutated to cause the highest possible disruption or change to the RNA structure without affecting neighbouring stem-loops, while keeping the same amino acid sequence and dinucleotide ratio (i.e., CpG and UpA). Predicted WT and mutated stem-loops visualised in Forna web server are shown. Nucleotides highlighted in green represent mutated positions, while red brackets represent positions of the hairpin loop in the WT structures and their altered position in the disrupted structures after mutagenesis.

**Figure 7.**
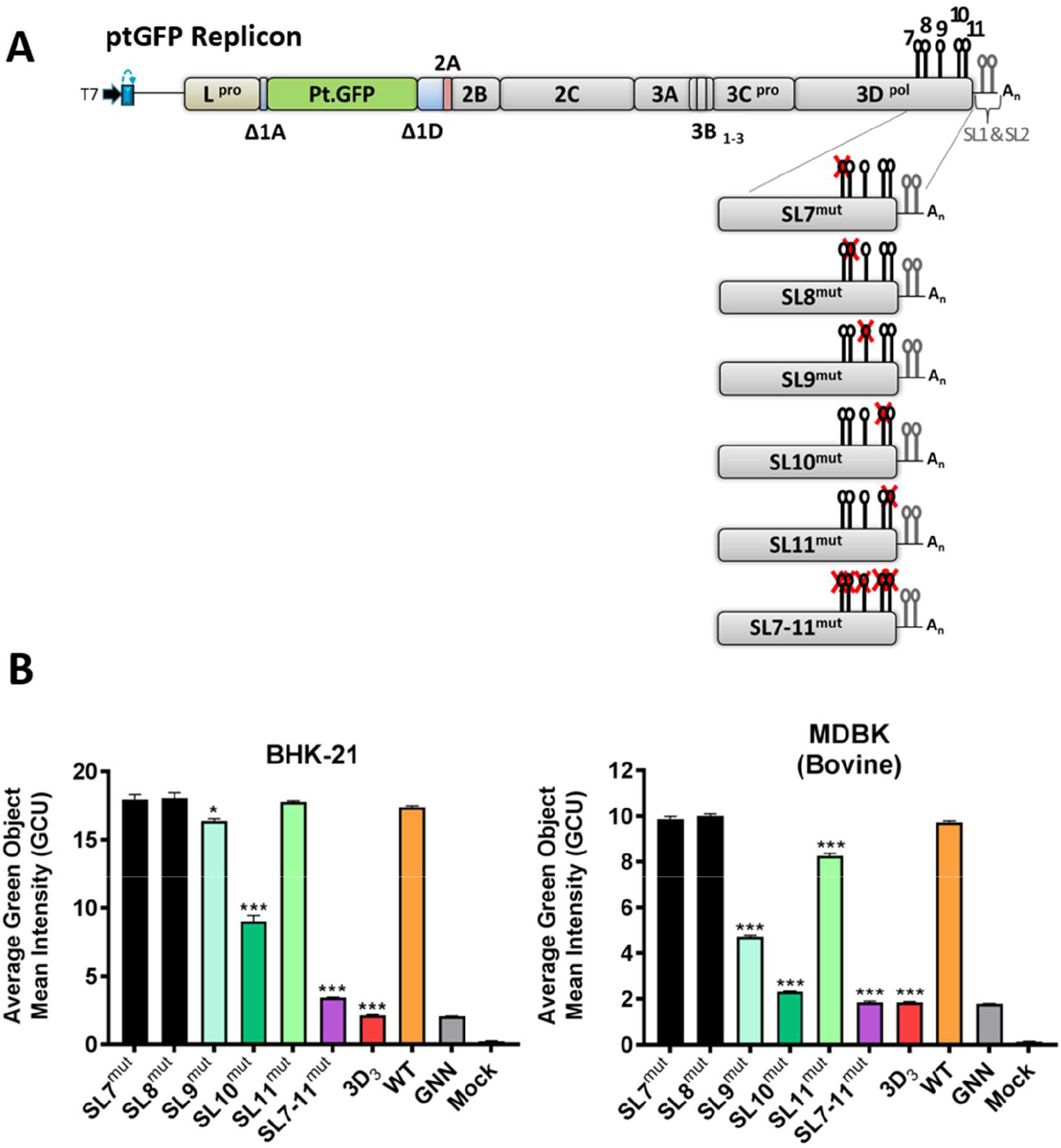
Effect of individual stem-loop (SL7 - SL11) mutagenesis on replication of the FMDV replicon. **(A)** Schematic representation of FMDV replicon constructs containing stem-loop mutations (SL9^mut^ – SL11^mut^). Sequence inserts containing stem-loop mutations were cloned directly into the ptGFP-replicon using the unique restriction enzymes BamHI and BspEI. **(B)** IncuCyte data represent the average cell (green object) GFP intensity per well at 8 h post-transfection within BHK-21 and MDBK cells. Results are the mean of three independent experiments ± standard error. Significant differences between WT ptGFP and SL^mut^ replicons were determined (*, P < 0.05; ***, P < 0.001).

To investigate whether the combined mutagenesis of SL9, SL10 and SL11 has a detrimental effect on replication of the FMDV replicon, constructs with two loops disrupted (SL9,10^mut^ and SL9,11^mut^), or all three loops disrupted (SL9-11^mut^) were tested as described above for the individual stem-loop mutations (Fig. 8A). In both cell lines, disruption of SL9 in combination with SL10 (SL9,10^mut^) resulted in a marked reduction of replicon replication when compared to replicons with the SL9 and SL10 mutated individually (see Fig. 7B and 8B). Replication of the SL9,10^mut^ replicon was severely disrupted (GFP intensity equal 27% (p-value < 0.001) and GFP intensity equal 20% (p-value < 0.001) of the GFP signal of the WT replicon in BHK-21 and MDBK cells, respectively), with replication levels comparable to the SL7-11^mut^ negative control (Fig. 8B). Interestingly, disruption of SL11 in combination with SL9 (SL9,11^mut^) resulted in a significant reduction of replicon replication in both cell lines (GFP intensity equal 60% (p-value < 0.001) and GFP intensity equal 36% (p-value < 0.001) of the GFP signal of the WT replicon in BHK-21 and MDBK cells, respectively; Fig. 8), although in BHK-21 cells individual mutation of SL9 and SL11 had only a marginal or no effect, respectively (see Fig. 7). Our computational prediction of SL9,11^mut^ did not suggest any disruption of the SL10 secondary structure, which is indirectly confirmed by the experimental data where replication impairment caused by joint permutation within SL9,11^mut^ is significantly less than that of the SL9-11^mut^ (GFP intensity equal 60% (p-value < 0.001) vs GFP intensity equal 28% (p-value < 0.001) of the GFP signal of the WT replicon in BHK cells, and GFP intensity equal 36% (p-value < 0.001) vs GFP intensity equal 20% (p-value <0.001) of the GFP signal of the WT replicon in MDBK cells, Fig. 8B). In both cell lines tested, disruption of all three stem-loops (SL9-11^mut^) resulted in a replication profile comparable to the SL7-11^mut^ (Fig. 8B). Table 3 summarises effect of mutagenesis of each of these stem-loops (individually and in combination) on replication of the FMDV replicon.

**Table 3.**
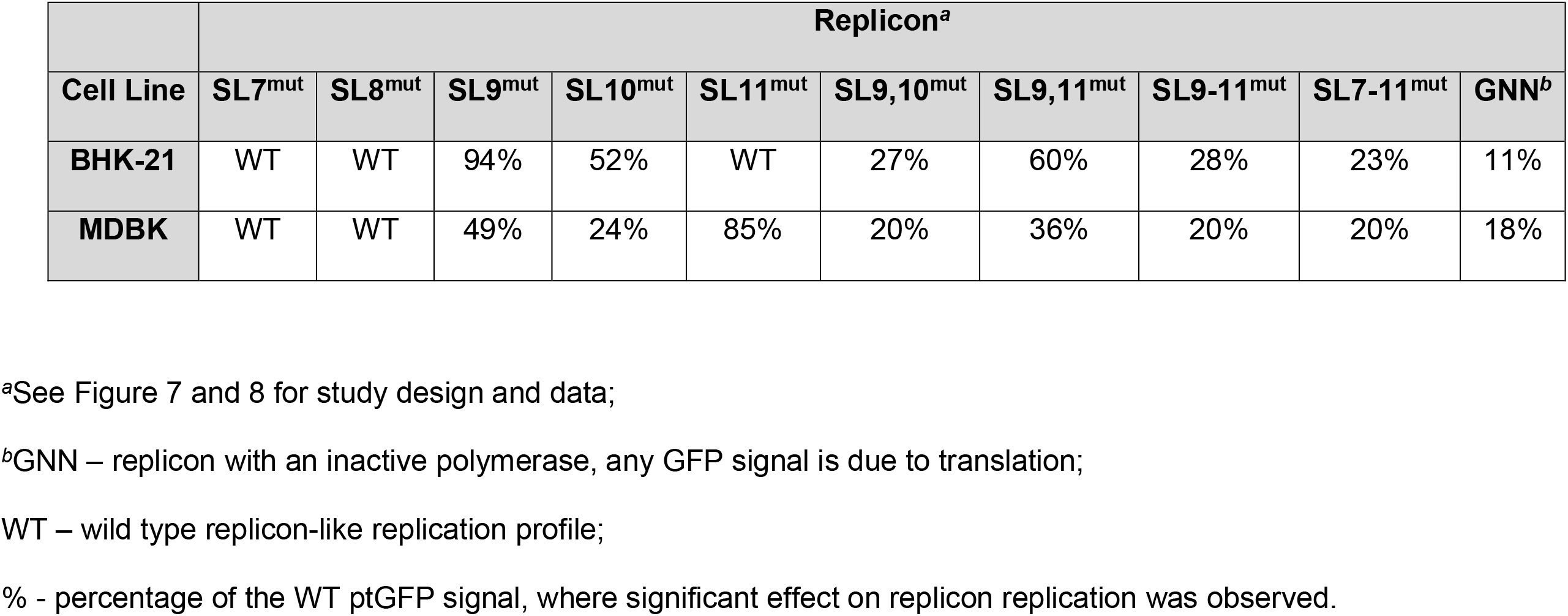
Summary of replication profiles of FMDV replicons after mutagenesis of conserved stem-loops localised within the 3D_3_ genomic region

**Figure 8.**
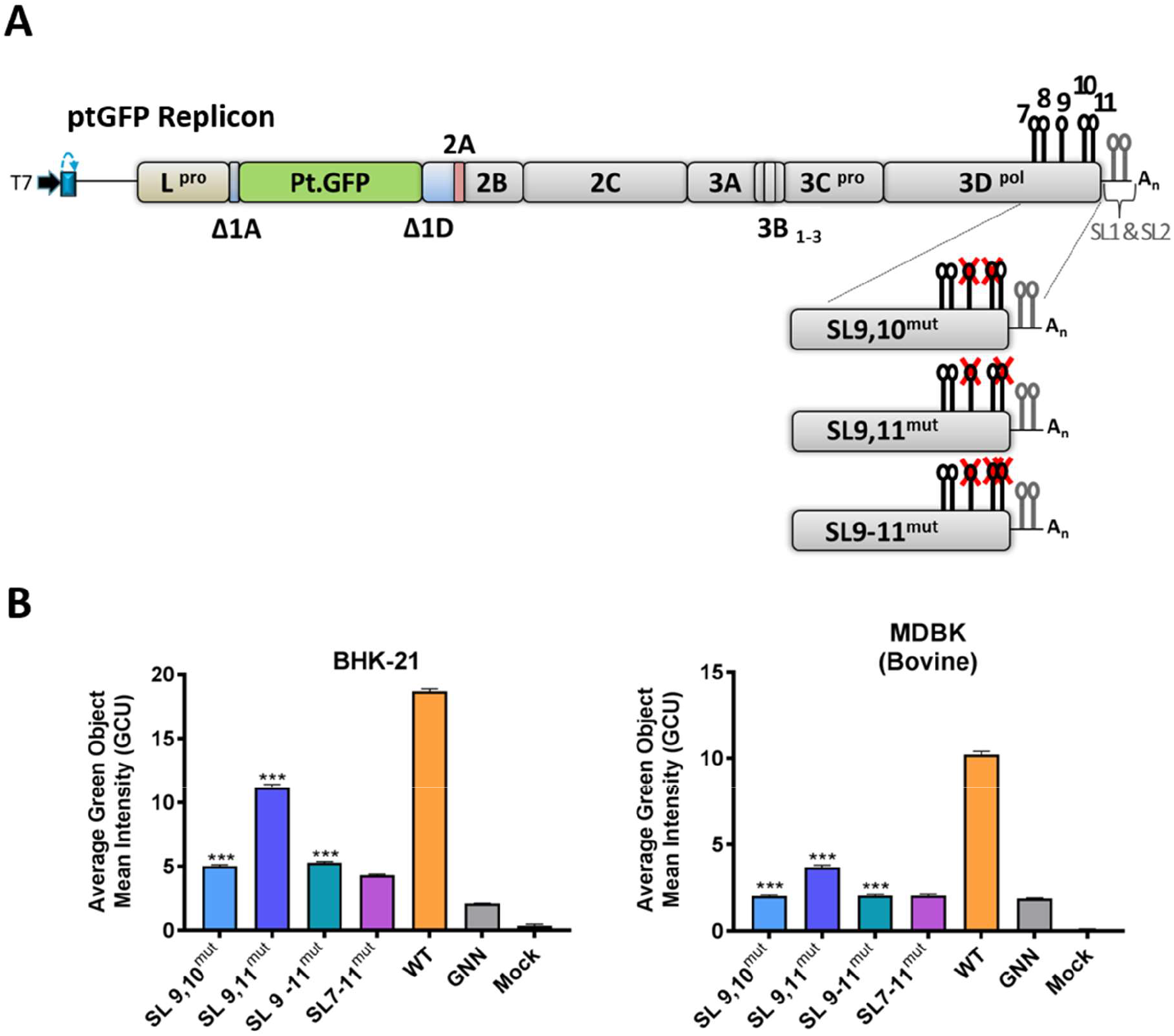
Effect of combined mutagenesis of stem-loops 9, 10 and 11 on replication of the FMDV replicon. **(A)** Schematic representation of FMDV replicon constructs containing combined stem-loop mutations (SL9,10^mut^, SL9,11^mut^ and SL9-11^mut^). Sequence inserts containing stem-loop mutations were cloned directly into the ptGFP-replicon using the unique restriction enzymes BamHI and BspEI. **(B)** IncuCyte data represent the average cell (green object) GFP intensity per well at 8 h post-transfection. Results are the mean of three independent experiments ± standard error. Significant differences between WT ptGFP and SL^mut^ replicons were determined (***, P < 0.001).

### Comparison of the conserved stem-loops within the FMDV 3D_3_ region to structures found in the 3’ terminal 3D encoding region of poliovirus

Two stem loops (referred to as loop α and β in Song *et al.* 2012) necessary for poliovirus (PV) replication are present in the 3’ terminal encoding sequence of PV 3D^pol^ (37, 38). Since PV is a member of a different genus in the family *Picornaviridae* and distantly related to FMDV, we investigated whether any of the stem-loop structures found in the 3’ end of the 3D^pol^ encoding region of the FMDV genome were similar to those present in the equivalent part of the PV genome. Therefore, we compared each of the FMDV RNA structures (SL7 to SL11) to the PV loops α and β using RNAforester. As described in Table 4, the structures identified in the 3’ terminal part of the coding region of FMDV 3D^pol^ do not appear to resemble those found in the equivalent position of the PV genome, while (using the same approach) the *cre* structures of PV and FMDV showed some structural similarity.

**Table 4.** Similarity^*a*^ comparison of RNA structures within the 3D^pol^ encoding region of FMDV and PV, calculated using RNAforester program

| **PV \ FMDV** | **SL7** | **SL8** | **SL9** | **SL10** | **SL11** | ***cre*<sup>\*</sup>** |
| --- | --- | --- | --- | --- | --- | --- |
| **$\alpha$** | -1.46 | -1.83 | -2.30 | -2.10 | -1.85 | nd |
| **$\beta$** | -1.18 | -2.07 | -2.52 | -2.27 | -2.07 | nd |
| ***cre*<sup>\*</sup>** | nd | nd | nd | nd | nd | 0.30 |

<sup>a</sup>Relative similarity scores equal to 1 are for two identical structures: the greater the distance from 1, the less structure similarity between two compared features. For simplicity, the output of RNAforester was rounded up to two decimal places.
<sup>\*</sup>comparison of *cre* of PV to *cre* of FMDV acts as a control of the structure prediction and RNAforester analysis.

## DISCUSSION

Many aspects of FMDV replication remain poorly understood, such as the function of RNA structures found within the ORF. Here we revisited the RNA structural architecture of the FMDV genome and, for the first time, investigated whether the putative stem-loops localised within the ORF are required for viral genome replication. Our results are in line with previous studies showing that FMDV has extensive RNA structure throughout the genome, substantially exceeding that found in viruses of other genera of the family *Picornaviridae* (e.g. MFED value >10% for FMDV genomic sequences comparing to <4% for viral sequences belonging to genus *Enterovirus*, *Hepatovirus*, *Parechovirus* and *Teschovirus*) (12, 56, 59). When compared to the previous structure predictions performed by Witwer *et al.* 2001, our study identified a greater number of conserved RNA structures within the FMDV ORF (53 stem-loops, with some merging into 46 branched structures, versus 25 structures predicted previously). Since we used a larger dataset than the previous authors (118 relatively diverse FMDV sequences versus nine used by Witwer *et al.* 2001), it is possible that we obtained a stronger statistical signal supporting conservation of these additional structures. Importantly, we found that three of the structures within the coding region of 3D^pol^ (i.e., SL9, SL10 and SL11) are critical for efficient replication of an FMDV replicon, thereby implying that they would provide the same function during virus replication.

Despite consistently elevated MFED values, the FMDV capsid encoding region contained only four RNA stem-loops which were conserved in all serotypes. Viral genomes characterised by high MFED values and low conservation of individual RNA structures have been observed before (66). For instance, the coding region of hepatitis C virus (HCV) showed elevated MFED values, while, except for the terminal genomic regions, the individual stem-loop structures were distinct between different HCV genotypes and even subtypes (66–68). Similarly, FMDV showed dense serotype-specific RNA structure within its capsid encoding region, which were not shared among other serotypes (as found in (66) and independently in here).

To identify functional RNA structures, we applied the CDLR algorithm to permute a genomic FMDV sequence (60). While the degree of possible mutagenesis is necessarily limited by protein coding, dinucleotide frequency, and codon usage constraints, the CDLR algorithm substantially disrupted secondary RNA structure of the native FMDV sequence in all regions apart from region encoding for 2C (Table 2 and Fig. S2). Since the permutation of the entire Δ1D-3A encoding region (which resulted in more extensive changes to the RNA structure) had a minimal effect on replication of the FMDV replicon, it is safe to state that conserved RNA stem-loops within the 2C encoding region are not essential for replicon of the FMDV replication *in vitro*.

Contrastingly, the CDLR scanning method identified three structures located at the 3’ terminal part of the 3D^pol^ encoding region that were important for replication of the FMDV replicon. Of these, SL10 showed the highest degree of pairing conservation and appeared to be the predominant structure important for replication of the FMDV replicon. Mutation of SL9, SL10 or SL11 showed a much greater reduction of replicon replication in MDBK cells compared to BHK cells. MDBK cells have been shown to secrete high levels of interferon (IFN) upon stimulation (69), while BHK-21 cells are known to lack an intact IFN pathway (70, 71). Furthermore, a number of published results suggest that RNA structure might directly or indirectly play a role in the modulation of antiviral responses (42, 54, 72, 73). Collectively, these observations suggest that SL9, SL10 and SL11 could play additional roles in the evasion of antiviral responses, and therefore mutation of these structures led to a drastic reduction in replication of the FMDV replicon in IFN-competent cell lines. In both cell lines tested, deletion of two or more stem-loops (SL9, SL10 and SL11) in combination significantly impaired replication of the replicon, suggesting that even in the absence of a fully functional antiviral pathway all three stem-loops are important for FMDV replication. Similarly to the viral genome, replication of an FMDV replicon involves viral protein synthesis, and the sequential synthesis of negative-(i.e. complementary) and positive-strand (i.e. genomic) viral RNA. Thus, although SL9-11 are required for replication of the replicon further studies are required to dissect which of these process (viral RNA translation and/or viral RNA replication) are dependent on SL9, SL10 and SL11. Interestingly, in the PV genome stem-loops within the coding region of 3D^pol^ have been identified that are requited for viral RNA synthesis (37, 38). However, these structures do not appear to share sequence or structural similarity with SL9, SL10 or SL11 in the FMDV genome.

The observation that replication of the FMDV replicon mutants with disrupted RNA structure elsewhere in the regions encoding the nsps (i.e., spanning 1D through to most of 3D^pol^) was surprising. The maintenance of extensive conserved internal base-pairing and consistently elevated MFED values observed in the relatively diverse set of FMDV isolate sequences analysed indeed strongly argues that the RNA structures formed by those genomic regions must play some functional role in the FMDV replication cycle. It is possible that at least some of the apparently ‘non-functional’ RNA structures are genome-scale ordered RNA structure (GORS) which may play a role in persistence of FMDV in its natural host (56, 59). While FMDV causes an acute disease in domestic animals (14, 74), it is known to persist in African Buffalo (*Syncerus caffer*), which are a natural reservoir of the virus (75–78). Since FMDV and African Buffalo are thought to have co-evolved together, it is possible that GORS developed in the FMDV genome as a part of the virus-host co-adaptation, where they might assist in evasion of immune recognition. The link between GORS, persistence and ability to minimise antiviral sensing has been shown for number of unrelated viruses (56, 59, 66, 72). Work is currently underway to investigate whether any of these remaining structures play a role in modulation of the antiviral sensing during FMDV replication in its natural host environment.

Although the function of the apparently non-essential RNA structures within the regions encoding the nsps remains to be defined, due to their conserved nature, they form a potential target for genome-scale attenuation of a wide range of FMDV strains. Such a strategy could contribute to the development of live attenuated FMD vaccines that may improve on the short duration of immunity, which is a shortcoming of current inactivated vaccines. Alternatively, the manipulation of RNA structures such as SL9, to provide attenuation in bovine cells but retain efficient growth in vaccine production cell lines (BHK), could be used to enhance biosafety of inactivated vaccine production. The hazards associated with the large-scale production of killed vaccine viruses include both accidental release of virus from high containment production facilities, and the distribution and use of improperly inactivated FMD vaccines (79–81).

In summary, we have generated a comprehensive map of RNA secondary structure located within the ORF of the FMDV genome and identified novel stem-loops within the coding region for 3D^pol^ that appear critical for FMDV replication. While the function of the other conserved structures remains to be determined they can be targeted to improve understanding of the FMDV biology. In addition, they have the potential to help develop safer FMDV vaccines, an idea which has been proposed for other viruses (6, 56, 82). We also show that usage of the CDLR algorithm can be successfully utilised to permute RNA sequences in search of functional RNA structures, which can be applied beyond viral RNA molecules using a freely available and easy to use package (60).

## MATERIALS AND METHODS

### Cell lines

Madin-Darby bovine kidney (MDBK) and baby hamster kidney (BHK-21) cells were obtained from the American Type Culture Collection (ATCC) and maintained in Dulbecco’s Modified Eagle Medium containing either 10% foetal bovine serum (FBS) or 10% horse serum (MDBK cells) at 37 °C and 5% CO_2_.

### FMDV sequence dataset

Full genome sequences of 105 viruses were selected from GenBank database based on nucleotide distance of their 1D encoding region, ensuring that they represent sequence variability that is known to be present between all seven FMDV serotypes (Table 5). Since sequences of SAT serotypes are the least represented on public databases, 13 additional full genome sequences of field SAT isolates (SAT 1 = 2, SAT 2 = 4 and SAT 3 = 7) were generated for the purpose of this study (isolates: SAT1/TAN/3/80, SAT1/ZAM/2/88, SAT2/BOT-BUFF/7/72, SAT2/MOZ/1/70, SAT2/ZAM-BUFF/18/74, SAT2/ZIM/8/89, SAT3/BOT/209/67, SAT3/RHO/26/76, SAT3/RHO/3/75, SAT3/SAR/9/79, SAT3/ZAM/P2/96(MUL-4), SAT3/ZIM/P25/91(UR-7), SAT3/ZIM/P26/90(HV-5) using methodology previously described (45).

**Table 5.** FMDV sequences selected from GenBank

| **Serotype:** | **Number of isolates:** | **GenBank accession numbers:** |
| --- | --- | --- |
| A | 19 | AY593788, MH053305, JF749843, HM854024, HQ832580, MH053306, KM268896, AY593802, KJ608371, MH053307, AY593751, AY593754, AY593761, AY593764, AY593766, AY593767, HM854022, AY593791, AY593794 |
| Asia 1 | 12 | AY593795, AY687334, DQ533483, DQ989306, DQ989315, DQ989319, EF149010, EF614458, HQ632774, JF739177, KM268898, MF782478 |
| C | 6 | MH053308, KM268897, MH053309, AJ133357, MH053310, AJ007347 |
| O | 21 | AY593819, MH053313, MH053311, MH053312, KF112885, KJ206909, HQ632769, HQ632771, KU291242, KR401154, GU384683, KF694737, AJ539140, MH053315, JX040491, MH053317, MH053318, MH053316, KJ560291, DQ404170, KU821591 |
| SAT 1 | 19 | AY593838, AY593845, MH053319, AY593844, JF749860, MH053321, AY593846, AY593839, AY593842, AY593841, AY593840, MH053322, AY593843, KM268899, MH053323, MH053324, MH053325, MH053326, MH053327 |
| SAT2 | 15 | MH053330, MH053332, MH053328, MH053329, JX014255, MH053333, AY593849, JX014256, AY593847, MH053335, KM268900, JF749862, MH053336, MH053337, KU821592 |
| SAT3 | 13 | AY593853, AY593851, MH053339, MH053340, MH053344, MH053343, AY593850, KJ820999, MH053341, MH053351, KX375417, KM268901, MH053350 |

### Prediction of conserved RNA structures within the FMDV genome

The genomic sequences of the 118 FMDV field isolates were aligned using the MAFFT X-INS-i algorithm which, in addition to nucleotide identity, takes into account RNA secondary structure information (83, 84). This approach minimized the potential to overlook conserved RNA structures that might be hidden in a nucleotide alignment containing distantly related FMDV sequences. The multiple sequence alignment (MSA) was analysed using the RNAalifold program implemented in The ViennaRNA Package (61), using the following options: a ribosum scoring matrix, calculating the partition function and base pairing probability matrix in addition to the minimum free energy (MFE) structure, producing structure without lonely pairs and with dangling energies added for the bases adjacent to a helix on both sides. Then, the conserved RNA structures in the full genome were ‘tidied up’ by removing gaps and long-distance interactions (i.e., interactions which were separated by 400 nucleotides or more). The same was repeated for each FMDV serotype individually (using the dataset described above) and serotype-specific conserved RNA structure prediction was compared to the conserved structure prediction for all 118 FMDV sequences. Only stem-loops which were verified in all seven FMDV serotypes were considered as highly conserved. Finally, the conserved, whole genome FMDV RNA structure was visualised by drawing a dot plot graph using an awk script written *in house* and available upon request. To visualise shorter genomic fragments containing predicted conserved RNA structure(s) (e.g., the 3’ terminal part of the 3D^pol^ encoding region and individual loops) in more detail, a particular genomic region together with its conserved structure prediction was extracted and visualised using an on-line Forna tool implemented in The ViennaRNA Web Services (85). The extent of nucleotide conservation in sequence forming hairpin loops of RNA structures (Fig. 5) was visualised using WebLogo 3.7.4 web server (86, 87).

Pairwise distance and MFED for full genome sequences of all seven FMDV serotypes (dataset described above) were prepared using the Sequence Distances and Folding Energy Scan programs implemented in SSE v1.4 package (60), respectively. The MSA for MFED analysis was prepared as described above, while controls for calculation of MFED were generated by the NDR algorithm. For sequence distance analysis the FMDV genomes were separated into three genomic regions: the 5’ UTR, the ORF and the 3’ UTR which were aligned individually by different MAFFT algorithms. The 5’ and 3’ UTRs were aligned by MAFFT X-INS-i, while the nucleotide sequence of the ORF was firstly converted into amino acid sequence using TRANSEQ EMBOS program (88), aligned using MAFFT G-INS-i (89) and then such generated amino acid alignment was converted into nucleotide sequence using TRANALIGN EMBOS program (88). All aligned genomic fragments were manually combined into a single MSA containing FMDV whole genomes. For both analyses the mean values for successive 400 base fragments with 20 nucleotide increment across the genome were plotted.

The average MFED values of the regions encoding the nsps of the FMDV isolates (i.e., dataset described above), the ptGFP-replicon and 50 CDLR-permuted ptGFP mutants were calculated as described above.

Since there appears to be a lot of ambiguity around the poly(C) tract, that region and its flanking positions were excluded from all analyses.

### *In silico* design of mutants containing modified segments within the non-structural encoding region

The regions encoding the nsps of the FMDV genome were chosen for mutagenesis by restriction site usage (sequence listed in Fig. 3A-B). To disrupt RNA secondary structures predicted in each restriction fragment of native FMDV genomes, sequences were mutated using the CDLR algorithm implemented in the Scramble Sequences Program of the SSE v1.4 package.

Structure prediction of the 3’ terminal part of the 3D^pol^ encoding region of the WT replicon which was scrambled by the CDLR algorithm (the 3D_3_ region) was generated as described above but using RNAfold (61), and using parameters corresponding to the ones applied in RNAalifold. The predicted structure was visualised in Forna.

To ‘quantify’ the difference between structure of the WT and scrambled replicons (Fig. 3A-B, Table 2), the whole genomic sequence of WT and each scrambled replicon was predicted using RNAfold, and fragments of the RNA secondary structure prediction corresponding to the permutated regions encoding the nsps (Fig. 3A-B) were compared using RNAforester and global alignment, with the relative scores as a measure of structure similarity (61, 63, 64).

For each predicted RNA structure located at the 3’ terminal part of the FMDV 3D^pol^ encoding region (SL7 - SL11 in the 3D_3_ region) nucleotides were changed manually (see results section). Individual putative stem-loops and their mutants were predicted using RNAfold implemented in The ViennaRNA package and mfold RNA structure prediction server (90), and were visualised using Forna RNA secondary structure visualisation tool.

### Comparison of putative RNA structures located within 3’ terminal 3D^pol^ encoding region of FMDV and PV

Computational prediction of two conserved PV RNA structures located in the 3’ terminal 3D^pol^ encoding region (termed loop α and β as in Song *et. al.* 2012) and described previously (37, 38) was repeated as described above. This was done as there was some discrepancy between the two publications about the exact structure of the two PV stem-loops. PV sequences representing variability of the PV 3D encoding region (GenBank accession numbers: NC_002058.3, DQ890388.1, FJ769378.1, EU794963.1, AY560657.1, HF913427.1, EU794957.1, EU794956.1, AF538842.1, EU684057.1, AF405667.1, AF405666.1, KJ170457.1, KJ170438.1, KU866422.1, AM884184.1, AJ132961.1, MG212491.1, MG212488.1, MG212485.1, MG212463.1, MG212456.1, MG212441.1, MG212440.1, KY941933.1, KY941932.1, KR259355.1, KC784372.1, KC880377.1, JX275352.1, JX274995.1, KX162704.1) were used. The RNA loop α and β were isolated and their structure aligned to the 3’ terminal part of the 3D^pol^ encoding region containing FMDV stem-loops SL7 - SL11 (3D_3_ region) using the RNAforester software and ‘small-in-large similarity’ calculation to determine whether any of the previously described PV stem-loops were similar to any of the FMDV RNA structures identified in this study. For more detailed analysis, each isolated FMDV putative RNA stem-loop (SL7 - SL11) was directly compared to PV loop α or β using RNAforester as described above.

### Clone construction

Sequences with mutations generated by the CDLR algorithm and nucleotide fragments containing mutated loops SL7 – SL11 were synthesised by custom DNA synthesis (GeneArt, Life Technologies) and provided within standard cloning vectors. These sequences were firstly sub-cloned into the pSP72 vector (Promega) to provide the unique restriction enzyme sites for subsequent cloning into the WT ptGFP replicon (Fig. 3A; (65)).

### *In vitro* transcription

Replicon constructs (5 μg) were linearised with AscI (New England Biolabs) for 1 h at 37 °C and purified using the E.Z.N.A. ™ Gel Extraction Kit (Omega Bio-Tek). Linear replicon DNA (500 ng) was added to transcription reactions at a final volume of 100 μl containing the following: Transcription Optimised Buffer (Promega), 10 mM DTT (Promega), 100 U RNasin Ribonuclease Inhibitor (Promega), 40 U T7 RNA polymerase (Promega), 20 mM rNTP’s (Promega) and nuclease-free water. Reactions were incubated at 37 °C for 2 h and the resulting transcript integrity assessed by agarose gel electrophoresis. RNA yield was quantified using the Quantus ™ Fluorometer (Promega), according to the manufacturer’s instructions.

### Cell transfection

Approximately 20 h prior to transfection cells were seeded into 24 or 12 well plates at the appropriate cell seeding density to achieve ~ 80% confluency. The following day, media was removed and replaced with FluoroBrite^TM^ DMEM (Gibco) supplemented with 2% FBS and 4 mM glutamine. Replicon transcript RNA (0.5-1 μg) was transfected into triplicate or quadruplicate cell monolayers using Lipofectamine 2000 transfection reagent as per the manufacturer’s recommendation (Thermo Fisher Scientific).

### Live cell imaging

Live cell image analysis was performed using the IncuCyte ZOOM kinetic imaging system (Essen BioScience) as described previously (62). Images were captured hourly for a period of 24 h with green fluorescent protein intensity measured using the integrated IncuCyte ZOOM image processing software. Data are shown as the average cell (green object) GFP intensity per well at 8 h post-transfection (where expression is at the maximum level).

### Statistical analysis

Replicon mutants were compared to WT ptGFP using one-way analysis of variance (ANOVA). Differences between groups were considered to be significant at a *P* value of <0.05 (*), <0.01 (**) or <0.001 (***). Error bars represent standard error of the mean (S.E.M.) of multiple independent experiments. Statistical analyses were performed with GraphPad Prism 8.00 (GraphPad Software, San Diego, California USA, www.graphpad.com).

### Data availability

Full genome FMDV sequences generated as a part of this study were submitted to GenBank and are available as following accession numbers: MW355668 - MW355680.

## ACKNOWLEDGMENTS

We thank colleagues in the WRLFMD (Pirbright, UK) for providing the FMDV isolates used in this study. The Pirbright Institute receives grant-aided support from the Biotechnology and Biological Sciences Research Council (BBSRC) of the United Kingdom (projects BB/E/I/00007035, BB/E/I/00007036 and BBS/E/I/00007037) providing funds to cover the open access charges for this paper. This work was supported by funding from UK Department for Environment, Food and Rural Affairs (Defra research project SE2943) and BBSRC research grant BB/K003801/1.

## Author Contributions

Lidia Lasecka-Dykes, Paolo Ribeca and Peter Simmonds performed bioinformatic analyses; Fiona Tulloch, Garry A. Luke, Lidia Lasecka-Dykes and Sarah Gold carried out experimental work and analysed data; Nick J. Knowles, Jemma Wadsworth and Mehreen Azhar selected and isolated viruses; Lidia Lasecka-Dykes and Caroline F. Wright sequenced FMDV isolates and analysed sequencing data; Fiona Tulloch, Lidia Lasecka-Dykes, Terry Jackson, Tobias J. Tuthill, Martin D. Ryan, Peter Simmonds and Donald P. King conceived and designed the experiments; Martin D. Ryan, Terry Jackson, Tobias J. Tuthill and Donald P. King directed the study; Martin D. Ryan, Terry Jackson, Tobias J. Tuthill and Donald P. King acquisitioned the funding; Lidia Lasecka-Dykes, Fiona Tulloch and Peter Simmonds wrote the initial draft of the manuscript; all authors reviewed and edited the manuscript.

