## Supplementary Materials for "Mutagenesis Mapping of RNA Structures within the Foot-and-Mouth Disease Virus Genome Reveals Functional Elements Localised in the Polymerase (3D^pol^) Encoding Region"

#### Supplementary Table S1

Supplementary legend for Figure 4

| | Pairings | | |
|---|---|---|---|
| | Compatible[a] | | Incompatible |
| **Colour** | **Number of pairings** | **Number of pairing types** | **Number of pairings** |
| Dark red | 118 | 1 | 0 |
| Red | 117 | 1 | 1 |
| Pink | 116 | 1 | 2 |
| Orange | 118 | 2 | 0 |
| Yellow | 117 | 2 | 1 |
| Pale yellow | 116 | 2 | 2 |
| Dark green | 118 | 3 | 0 |
| Green | 117 | 3 | 1 |
| Pale green | 116 | 3 | 2 |
| Dark turquoise | 118 | 4 | 0 |
| Turquoise | 117 | 4 | 1 |
| Pale turquoise | 116 | 4 | 2 |
| Dark blue | 118 | 5 | 0 |
| Blue | 117 | 5 | 1 |
| Pale blue | 116 | 5 | 2 |
| Grey | ≤115 | ≤6 | ≥3 |

[a]Compatible pairings are G-C, C-G, A-U, U-A, G-U and U-G;

White = Unpaired nucleotides within loops;

○ = indicates nucleotide positions where a substitution resulted in an alternative compatible pair;

Grey bars indicate unstructured regions between the stem-loops (not drawn to scale).

### Supplementary Figure S1

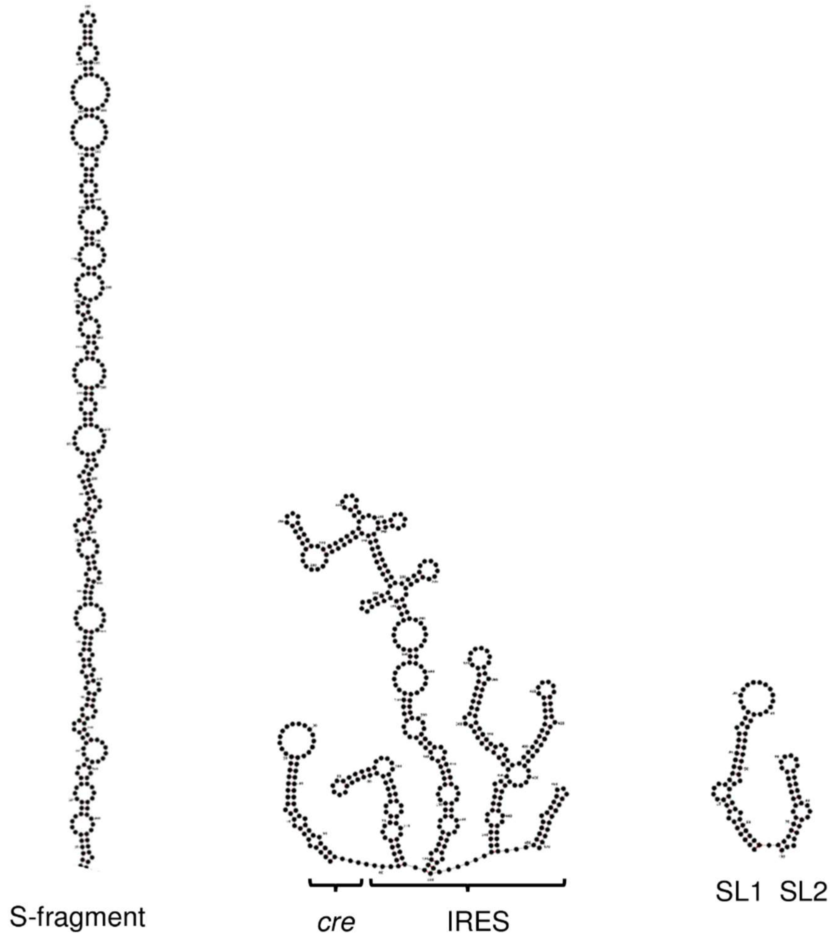


Conserved whole genome RNA structure was predicted for 118 FMDV filed isolates as described for Figure 1 and regions of previously published structures located within the 5' UTR (S-fragment, *cre* and IRES) or 3' UTR (SL1 and SL2) were visualised using Forna web service.

### Supplementary Figure S2

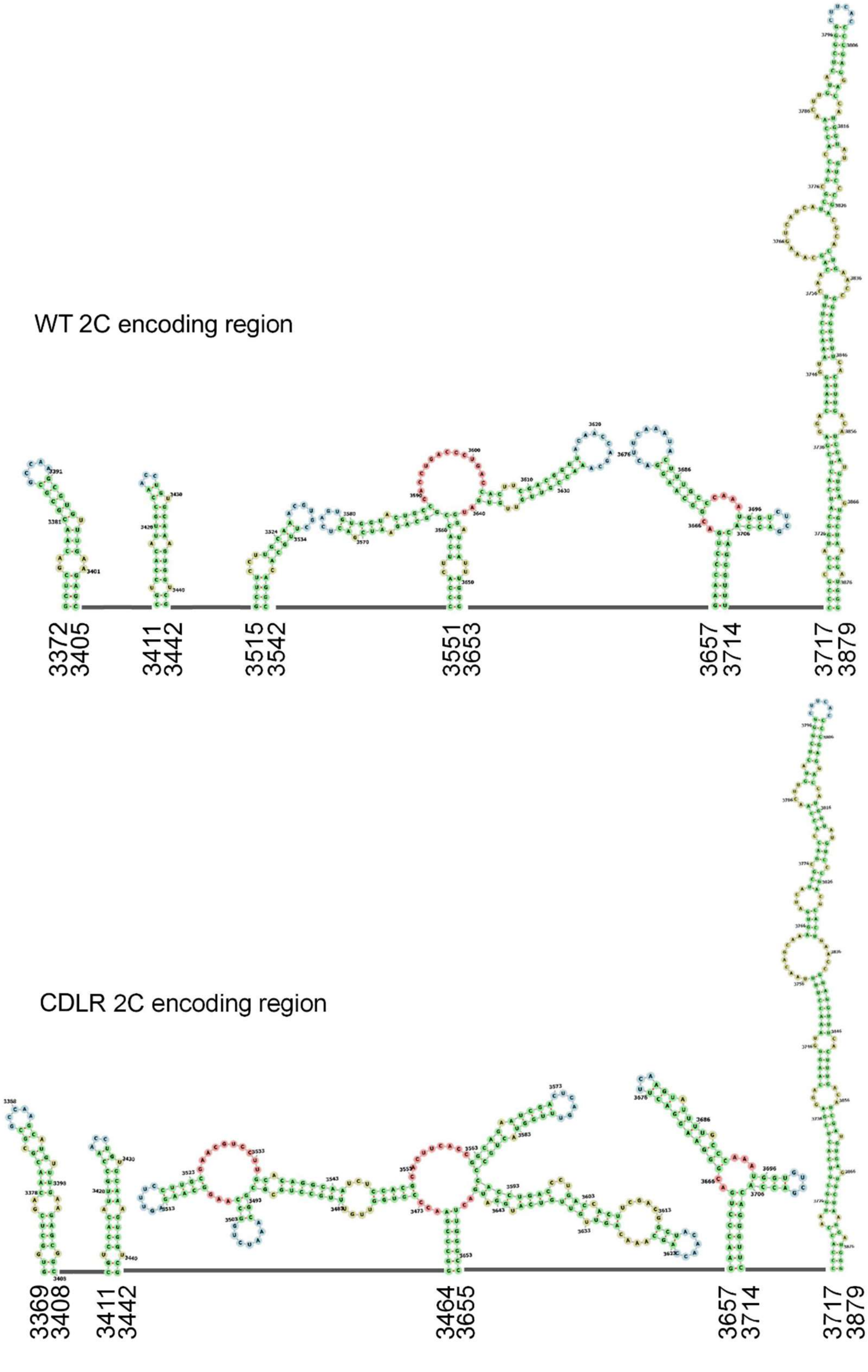

RNA structure for the whole genome of the WT replicon and a replicon with the 2C encoding region permuted by the CDLR algorithm was predicted using RNAfold. The predicted structure of the 2C encoding fragment was visualised using Forna web service. Lines represent unstructured regions and numbers at the beginning and the end of each individual stem-loop correspond to the genomic positions of each replicon. Nucleotides forming stems are shown in green, interior loops in yellow, junctions in red and hairpin loops in blue.

**Supplementary Figure S3**

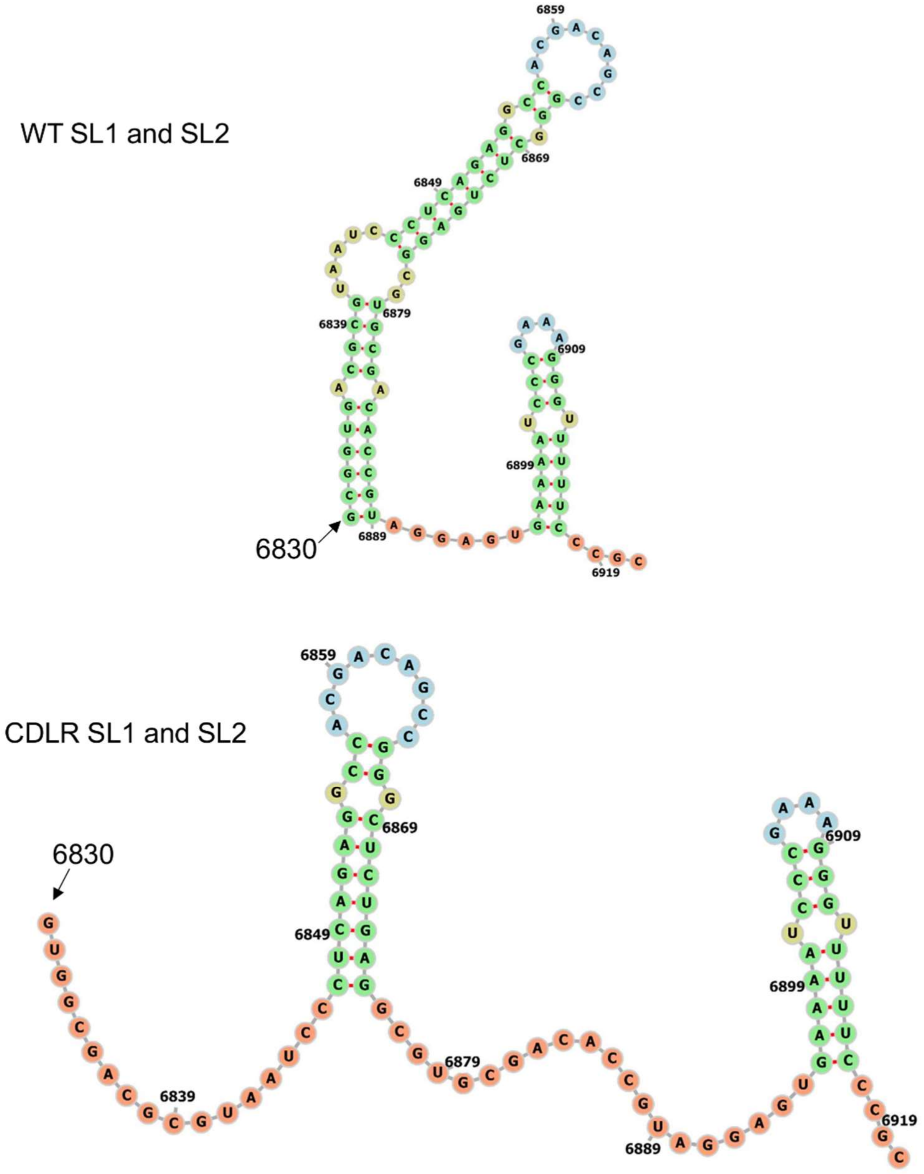


RNA structure for the whole genome of the WT replicon and a replicon with the $\Delta$1D-3D$_3$ encoding region permuted by the CDLR algorithm was predicted using RNAfold. The predicted structure of SL1 and SL2 stem-loops located in the 3' UTR was visualised using

Forna web service. Numbers correspond to the genomic positions of each replicon. Nucleotides forming stems are shown in green, interior loops in yellow, hairpin loops in blue and unpaired region in orange.

### Supplementary Figure S4

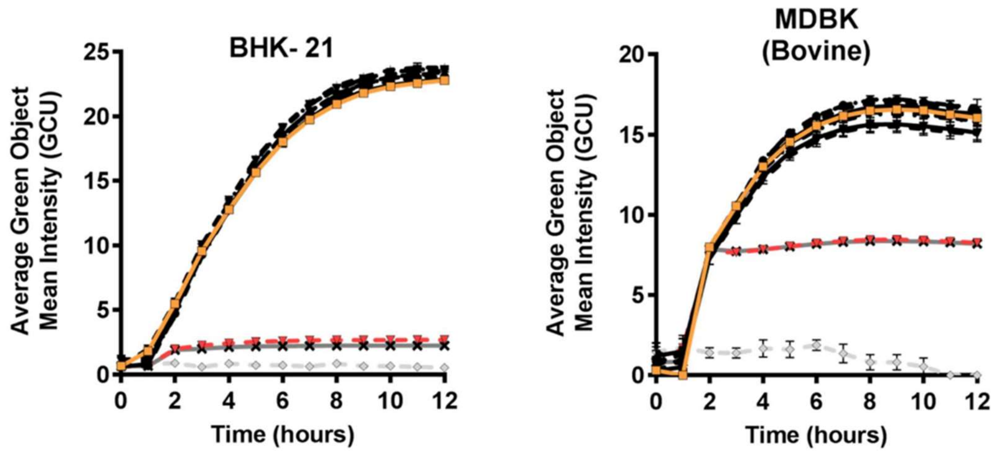


Replication kinetics of FMDV replicon constructs containing CDLR-permuted regions which were described in Figure 3. IncuCyte data represent the average cell (green object) GFP intensity per well over a period of 12 h. Replicon RNA from CDLR mutants (black (or red for $3D_3$)), WT ptGFP (orange) or ptGFP-GNN (grey) was introduced into BHK-21 or MDBK cell monolayers. Results are the mean of three independent experiments ± standard error.

### Supplementary Figure S5

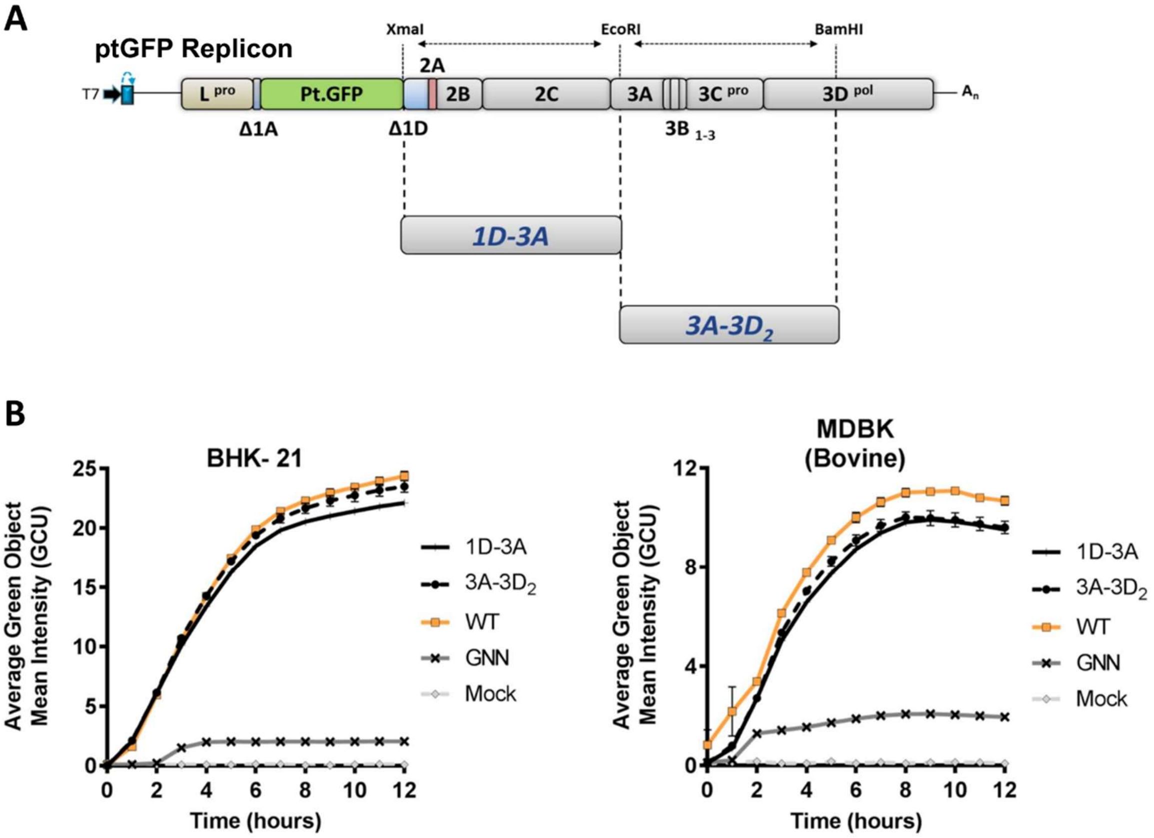


Replication kinetics of FMDV replicons containing larger sections of the genome permuted by the CDLR algorithm. **(A)** Schematic representation of CDLR replicons encoding larger mutated regions. Mutated inserts were cloned directly into the ptGFP-replicon using the unique restriction enzymes shown. **(B)** IncuCyte data represent the average cell (green object) GFP intensity per well over a period of 12 h within BHK-21 and MDBK cells. Results are the mean of three independent experiments ± standard error.

### Supplementary Figure S6

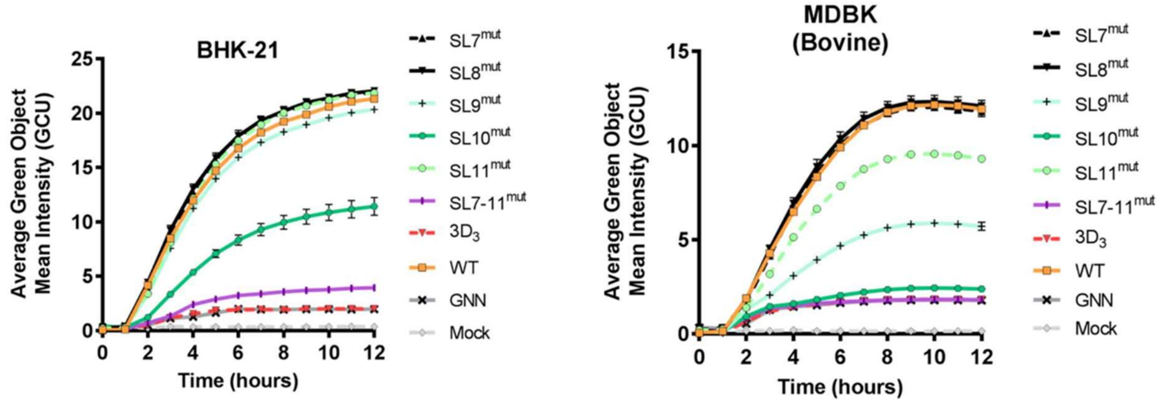


Replication kinetics of FMDV replicon constructs containing individual stem-loop mutants which were described in Figure 7. IncuCyte data represent the average cell (green object) GFP intensity per well over a period of 12 h within BHK-21 and MDBK cells. Results are the mean of three independent experiments ± standard error.

### Supplementary Figure S7

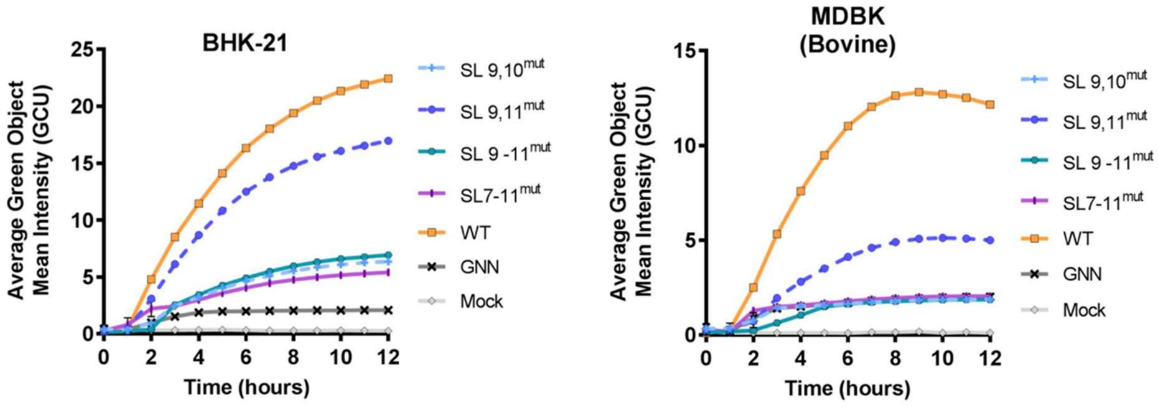


Replication kinetics of FMDV replicon constructs containing stem-loop mutants which were described in Figure 8. IncuCyte data represent the average cell (green object) GFP intensity per well over a period of 12 h within BHK-21 and MDBK cells. Results are the mean of three independent experiments ± standard error.
